# Spatial representation in CA1 superficial pyramidal cells is impaired after postnatal ablation of hippocampal Cajal-Retzius cells

**DOI:** 10.64898/2026.03.24.711525

**Authors:** Sachuriga, Kristian Moan, Keagan Dunville, Nina Seiffert, Ingvild L. Glærum, Giulia Fiori, Giulia Quattrocolo

## Abstract

Cajal-Retzius neurons (CRs) are a transient cell type that populates the postnatal hippocampus. To test how the persistence of CRs shapes the maturation of hippocampal function, we used a CRs-specific transgenic mouse line combined with targeted viral delivery to selectively ablate CRs in the postnatal hippocampus. Single cell sequencing revealed that gene networks in superficial CA1 pyramidal cells were more strongly perturbed compared to deep CA1 pyramidal cells. To test if these two subpopulations were also distinctly affected in their function, we performed in vivo recordings from spatially modulated cells in CA1. Our analysis showed an impaired spatial representation specifically in superficial CA1 pyramidal cells. Additionally, we observed an increased CA3 to CA1 excitatory drive, as indicated by increased gamma oscillations, and alterations of intrinsic firing properties in superficial CA1 pyramidal neurons confirmed by in vitro electrophysiological recordings. Together, these results indicate a crucial role for CRs in the maturation of hippocampal subcircuits.

## Introduction

Cajal-Retzius neurons (CRs) are transient cortical neurons known for their role in orchestrating cortical architecture during development ^1^, being the main source of Reelin at prenatal stages ^2–4^. Postnatally, CRs undergo programmed cell death and disappear from most cortices within the early postnatal weeks ^1^. However, we recently showed prolonged CRs survival in several brain regions, including association and prefrontal cortices, entorhinal cortex, and hippocampus ^5^. These regions exhibit delayed functional onset and protracted maturation relative to sensorimotor cortices ^6–9^. This intriguing spatiotemporal overlap between delayed functional maturation and sustained CR survival raises the question of whether CRs actively contribute to postnatal development within these specific brain regions.

We have previously shown that postnatal CRs are critical for organizing hippocampal networks^10^. In the absence of postnatal CRs, we observed significant layer-specific changes in cell morphology, dendritic spine profiles, and the expression of synapse-related genes and synaptic proteins ^10^. To study how postnatal CRs could contribute to the maturation of defined functional cell types, and the development of hippocampal circuit dynamics, we used an intersectional genetic approach ^5^ combined with neonatal viral injections to specifically ablate CRs from the postnatal hippocampus. Single nuclei sequencing of CA1 pyramidal cells revealed a greater alteration of gene networks related to plasticity in the superficial population of CA1 pyramidal cells, compared to the deep population. As in vitro physiological recording confirmed a stronger alteration in the firing properties of superficial than deep pyramidal cells, we tested if the function of the superficial population was also more specifically affected. Indeed, our in vivo recordings showed that postnatal ablation of CRs altered the upstream input arriving at CA1 and impaired spatial representations in superficial but not deep pyramidal cells. These findings demonstrate a critical role for CRs in the maturation of hippocampal circuits and suggest a differential influence on neuronal subpopulations.

## Results

### Targeting and Ablation of hippocampal CRs

In this study, we investigate whether prolonged survival of postnatal hippocampal CRs is crucial for hippocampal functional development. To define the role of postnatal CRs, we employed an intersectional genetic strategy by crossing Pde1c-Cre+/- mice^11^ with NDNF-flox-FlpO+/- mice to specifically target hippocampal CRs ^5^. In the resulting Pde1c-Cre+/-; NDNF-flox-FlpO+/- mice, we selectively ablated CRs postnatally using a Flp-dependent diphtheria toxin-expressing viral vector. A viral construct (pAAV-mCherry-flp-DTA) encoding pan-neuronal mCherry and Flp-dependent DTA was bilaterally injected into the dorsal hippocampus of postnatal day 0 (P0) pups. While all infected neurons expressed mCherry (Supp.Fig. 1A-B), DTA expression was restricted to Pde1c-Cre+/-; NDNF-flox-FlpO+/- cells, as NDNF- driven FlpO expression occurred only after Cre-mediated excision of the last coding exon, ensuring CRs-specific ablation. Immunostaining for CRs-specific markers p73 and Reelin (Supp.Fig. 1 C-F) at P60– 68 revealed an overall reduction of 82% in hippocampal CRs in Pde1c-Cre+/-; NDNF-flox-FlpO+/-; DTA (CR;DTA+) experimental animals compared to Pde1c-Cre-/-; NDNF-flox-FlpO-/-; DTA (CR;DTA-) controls (Suppl.Fig 1H). These findings, in line with our previous work, ^10^ validate the efficacy of this genetic ablation strategy for eliminating hippocampal CRs and provide a foundation for assessing their role in hippocampal function.

### Alterations in gene expression programs in superficial CA1 pyramidal neurons

Previous work has demonstrated that CRs distinctly regulate synaptic formation in the dendritic layers of CA1 ^10^. However, this work relied on bulk transcriptomic data and lacks resolution to identify cell-specific transcriptomic changes. To examine changes in gene expression at the single cell level in CA1 pyramidal cells, we performed single-nuclei RNA sequencing from dorsal hippocampus of control and experimental mice (11,619 nuclei from 4 mice; Supp. Fig. 2A) at P30. Nearest neighbors clustering identified major hippocampal cell classes, including CA1 pyramidal cells. CA1 cells were further divided into deep and superficial subtypes (Fig. 1A) by cross-referencing with the Allen Transcriptomic Brain Cell Mouse Atlas, 2020 single cell RNA seq data12. CA1 UMAP embeddings varied between the control and experimental samples (Fig. 1B). To understand how CA1 subclasses’ transcriptomic profile is altered by CR cell ablation, a total of 332 marker genes separating superficial and deep layer subtype were plotted (Fig. 1C), though marker gene expression appeared conserved across genotype. We performed DGE analysis within subtypes between control and experimental mice (Fig. 1D,E), and we found that few genes were differentially upregulated in either subtype (18 in deep, 8 shared, and 15 in superficial; Fig. 1D) and instead the largest effect was observed in downregulated genes in the superficial subtype (43 in deep, 85 shared, and 229 in superficial, Fig. 1E). Of those 398 genes, we then investigated how the top 100 differentially expressed genes (DEG) drive differences within pyramidal cell subpopulations between control and experimental mice. We found that genes were primarily downregulated when comparing the control to the CR;DTA+ subtypes and when we applied k-means clustering (k=4) across these DEG that clustered expression differences represented superficial control (cluster 2), CA1 control (cluster 4), deep control (cluster 3), and CA1 experimental mice (cluster 1, Supp.Fig. 2B). Among these 100 genes, we found several related to intrinsic firing properties and synaptic connections including Olfm1, Scn2b in cluster 3, Grina, Nrgn, and Caly in cluster 4, and Calm3, Calm1, and Junb in cluster 2; most genes, however, were related to ribosomal processing and cytoskeletal structure, suggesting an overall change in protein translation and in cell morphology, respectively, similar to previous observations in decreased expression of genes related to dendritomorphogenesis and synaptic structure 10. Overall, we observed a stronger transcriptomic response to CR-cell ablation in CA1 superficial cells compared to deep. However, each individual gene may contribute to a larger network or group of coexpressed genes that participate in a conserved biological process. To dissect genes with common regulatory networks, we relied on a recent framework13 to perform weighted correlation in gene networks of single cell subpopulations to relate function and connectivity. We performed high-definition weighted gene correlation network analysis (hdWGCNA14) to represent gene networks as pairwise gene co-expression to interrogate how gene networks differ between the control and experimental CA1 subtypes. We first performed our analysis in CA1 superficial cells in the CR;DTA- (Fig. 1F) and the CR;DTA+ mice (Fig. 1G) and found that the total number of de novo identified networks decreased from the control to the experimental mice. These results suggest transcriptional consolidation, in that cell-type specific gene networks became more exclusive to their associated subtype. However, when we performed differential module eigengene (DME) analysis, we discovered instead that aggregate gene expression of ∼14 networks were significantly changed in the CR;DTA+ superficial subpopulation compared to the control (Fig 1H). We identified 4 networks of interest as member genes were related to synaptic plasticity. Networks are denoted by their top module eigengene in Fig 1G as well as symbolically in following panels (Networks: Itm2b:spiral:S4; Mgat4c:star:S1; Syt7:square:S12; Cdh8:triangle:S13; Opcml:hexagon:S8[non-significantly changed]). In line with our previous results, we found that Itm2b network was significantly downregulated in the CR-cell ablated superficial cells. Itm2b encodes synaptic anchoring protein, BRI2, which has been shown to facilitate postsynaptic inputs to CA1 pyramidal cells15. Conversely, Opcml network (S8) was non-significantly upregulated though ubiquitously expressed in all neuronal cell types, corroborating a decline in cell-specific regulation of transcription. To further investigate *Itm2b* network, we queried the top 25 module eigengenes coexpressed in the network (Fig. 1I) and coexpressed genes were primarily related to synaptic maintenance (*Syp*, *C1qtnf4*, *Plxna4*), neuronal calcium modulation (*Camk1d*, *Calm2*, *Calr*, *Nrgn*), or transmission (*Grm1*, *Scg2*). Moreover, we found that *Grm1* is a selective marker in CA1 superficial cells, indicating that this network is specific to CA1 superficial cells. Using these networks, we queried gene ontology databases (GO:DB) and found that under the Molecular Function annotation, S4 network was significantly enriched for P-type Sodium transporter activity (Fig. 1J). P-type transporters are necessary for fast ATP-mediated membrane transport across synapses, and in the context of CA1, promote cell-cell connectivity and intrinsic firing properties ^16^. Additionally, S4 is enriched for postsynapse localization when queried in SynGO database (Fig. 1K). Other networks significantly changed in the DME analysis were found enriched in synaptic gene ontology including S1 for presynaptic dense core vesicle exocytosis, S12 for postsynaptic cytoskeleton, and S13 for anchored component of postsynaptic membrane (Fig. 1K). Overall, these results indicate a high degree of alterations to a wide range of networks related to specialized neuronal functions including plasticity and connectivity. Conversely, when we analyzed the deep CA1 cells in the CR;DTA- (Supp.Fig. 2C) and CR;DTA+ (Supp.Fig. 2D) animals we found a difference in manifold representation between both groups, however, only a slight decrease in overall network number. Additionally, DME analysis for deep CA1 subpopulation revealed that only 3 networks were upregulated (Supp.Fig. 2E), though none of the significantly altered networks appeared in the SynGO database (Supp.Fig. 2F), further suggesting minor perturbations to transcriptional regulation in the CA1 deep cells. Our transcriptional analysis revealed a high degree of differential expression of genes related to intrinsic activity and connectivity in CA1 cells. Moreover, analysis between subtypes highlighted significant yet selective perturbations in the CR;DTA+ superficial CA1 pyramidal cells. Thus, to assess potential deficits in CA1 pyramidal cell activity, we performed *in vitro* electrophysiological recordings alongside *in vivo* electrophysiological recordings in the adult CA1 of postnatal CRs ablated mice.

**Figure 1.**
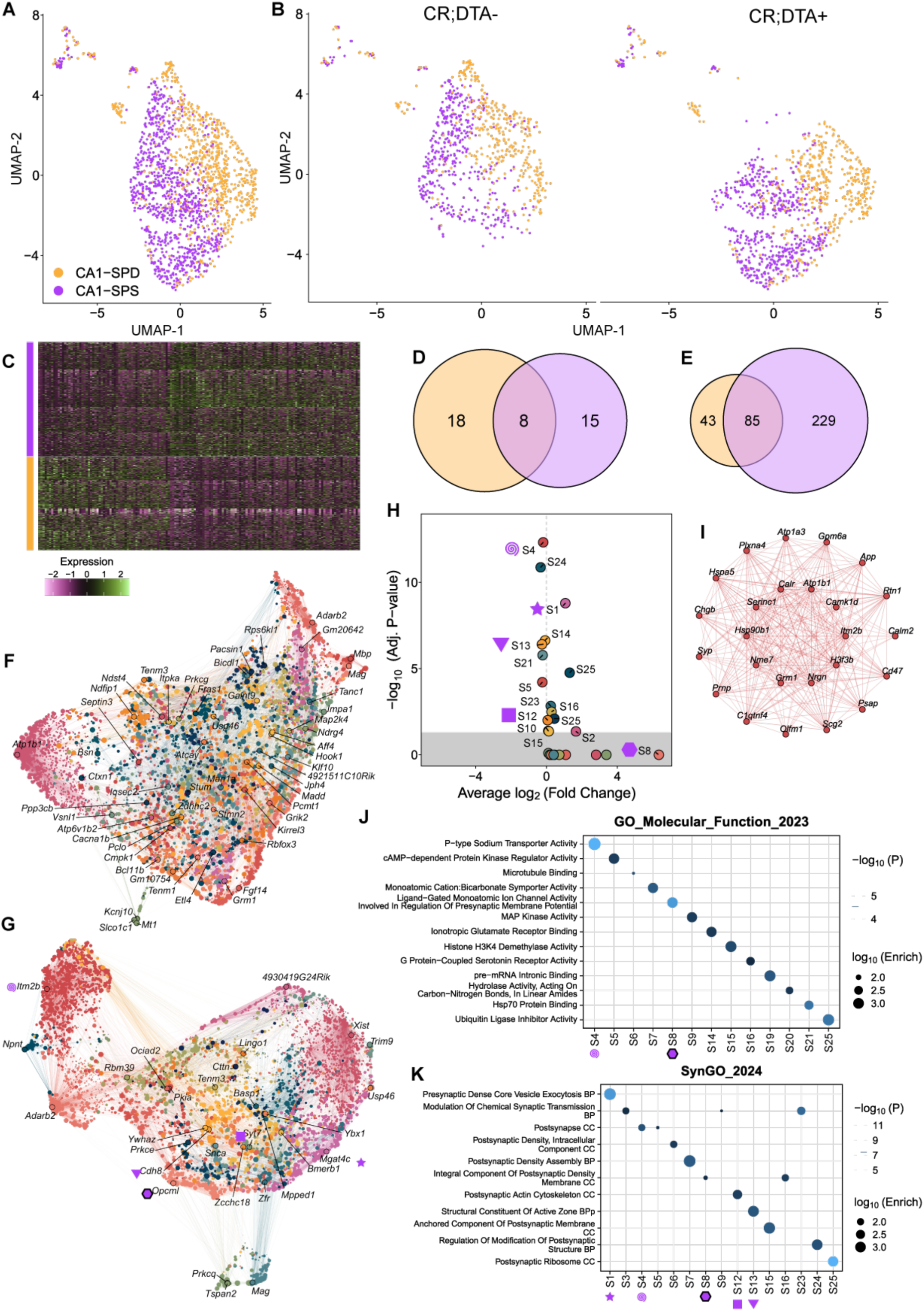
Cajal–Retzius cell ablation selectively remodels gene expression programs in superficial CA1 pyramidal neurons. (A) UMAP of subsetted CA1 cells, deep layer cells are labeled in yellow, superficial layer cells are labeled in violet. Cells were clustered using semi-supervised ANNOY method. (B) CA1 UMAP separated by experimental group. UMAP space occupation are dissimilar between superficial and deep layer cells in both the CR;DTA- and CR;DTA+. (C) Marker gene expression of ∼330 significantly up or downregulated genes. Gene expression is plotted from individual cells (∼2000 CA1 cells in total), cells under yellow bar are deep layer cells whereas cells under violet bar are superficial cells. Cut off for significantly regulated gene was set by adjusted p-value < 0.05 and |Log_2_FC| >= 0.25. (D) Upregulated gene overlap between deep (yellow) and superficial (violet) CA1 pyramidal cells. Differential gene expression was calculated either as *Gene Expression_CR;DTA+_ – Gene Expression_CR;DTA-_* for both deep and superficial cells. (E). Downregulated gene overlap between deep (yellow) and superficial (violet) CA1 pyramidal cells. Differential gene expression was calculated as *Gene Expression_CR;DTA+_ – Gene Expression_CR;DTA-_* for both deep and superficial cells. (F) Gene network UMAP for superficial cells in CR;DTA- CA1, demonstrating ∼52 networks of correlated gene expression. (G) Gene network UMAP for superficial cells in CR;DTA+ CA1, demonstrating ∼28 networks of correlated gene expression. Violet symbols indicate corresponding gene networks in the subpanels (H) Volcano plot of differential module eigengene network expression between CR;DTA- and CR;DTA+ CA1 superficial layer pyramidal cells. DME was calculated as *Aggregate Network Expression_CR;DTA+_ – Aggregate Network Expression_CR;DTA-_*. Cutoff set to adjusted p-val < 0.05 for significantly up or downregulated network. (I) Representative correlation plot of top 25 module eigengenes in network S4, the network represented by Itm2b in Figure 7H. Pairwise correlation values are represented by lines between MEs. (J) Gene Set Enrichment Analysis of Gene Ontology Database’s “Molecular Function” category. Only the top-scoring category is displayed per network. 13/28 networks were found to be significantly enriched, including S4 and S8. (K) Gene Set Enrichment Analysis of Synaptic Gene Ontology (SynGO) updated 2024 database. Only the top-scoring category is displayed per network. 15/28 networks were found to be significantly enriched, including S1, S4, S8, S12, and S13.

**Figure 2.**
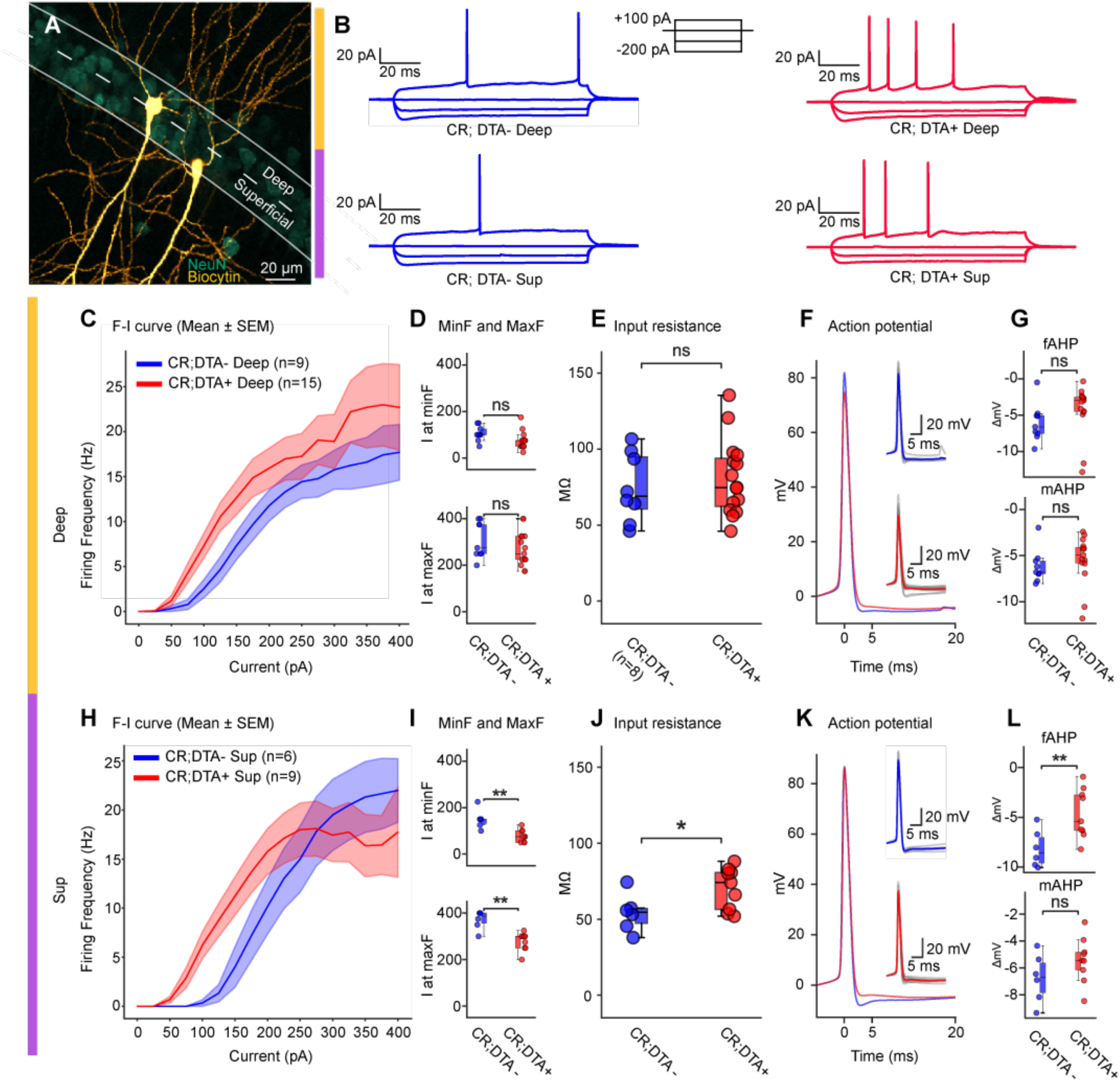
Selective enhancement of intrinsic excitability in superficial, but not deep, pyramidal neurons. (A) Representative image of deep and superficial CA1 cells from a CR;DTA+ animal. Scalebar bottom right. (B) One set of representative voltage traces from each cell group in these experiments. Four sweeps of current injections used to elicit voltage responses, illustrated in the top middle. (C) Frequency–current (F–I) curves for deep pyramidal cells from CR;DTA- (blue; n = 9 cells) and CR;DTA+ mice (red; n = 15 cells). Mean ± SEM firing frequency is plotted as a function of injected current (0–400 pA). (D) Current at minimal firing (Fmin) and maximal firing (Fmax) for deep cells; no significant differences between groups (ns). (E) Input resistance of deep CA1 pyramidal cells at a hyperpolarizing step of -50 pA; no significant difference. (F) Left: Comparison of average action potential from CR;DTA- (blue) and CR;DTA+ (red). All action potentials used for this test were extracted from the sweep which was +100 pA after the first sweep with at least 1 action potential. 0 on Y-axis represents start of action potential, which was defined as the point where the derivate of the voltage reached 20mV/ms. 0 on X-axis represents peak of action potential. fAHP was measured as minima between peak and 5 ms after peak. mAHP was measured from end of fAHP until 20 ms. Smaller illustrations show average action potential from both groups, in their respective color, and all individual action potentials in grey. Left: boxplots for comparison of fAHP and mAHP. 0 represents voltage at action potential start, while AHP amplitude is measured as difference between action potential start and the minimum voltage between 0 and 5 ms (fAHP) and 5 and 20 ms (mAHP). (G-J) Same as C-F, only for superficial CA1 pyramidal cells. * = p <0.05; ** = p <0.01

### Selective shift of intrinsic excitability in superficial, but not deep, pyramidal neurons

As our transcriptomic analysis suggested cell-type specific alterations, we questioned if intrinsic properties of the neurons could also be specifically differentially affected between deep and superficial pyramidal CA1 pyramidal cells. Thus, we performed whole-cell patch clamp experiments from the two subpopulations of CA1 pyramidal cells (Fig. 2A), in control and experimental animals. In response to +100 pA current injections, CR;DTA+ cells, especially superficial cells, appeared to be more excitable compared to their CR;DTA- counterpart (Fig. 2B). Measured changes in firing frequency in response to increasing depolarizing current steps showed that the frequency–current (F–I) relationship in deep pyramidal neurons (orange panels) was slightly shifted between CR;DTA- and CR;DTA+ (Fig. 2C). Still, the current needed to elicit firing and reach the maximal firing rate were not significantly different between groups (Fig. 2D). Similarly, input resistance did not differ between the deep cells groups (Fig. 2E). As the afterhyperpolarization phase is critical in regulating the frequency of firing^17^, we analyzed the amplitude of the fast and medium afterhyperpolarization after action potential firing and noticed that they were also unchanged (Fig. 2F-G). Together, these data indicate that CRs ablation has little effect on the basic excitability of deep pyramidal neurons.

In contrast, superficial pyramidal neurons showed several electrophysiological alterations following CRs ablation. F–I curves of superficial neurons were different between groups (Fig. 2H), and analysis of firing thresholds revealed significant changes in the current required to induce firing and to reach the maximum rate (Fig. 2I), indicating a modified excitability range in the CR;DTA+ group. Superficial neurons from CR;DTA+ animals also exhibited a higher input resistance compared with the CR;DTA- animals (Fig. 2J), suggesting the enhanced intrinsic excitability. Averaged action potential traces showed a less pronounced afterhyperpolarization (AHP) in CR;DTA+ cells (Fig. 2K), and quantification confirmed a significant decrease in fast AHP (fAHP) amplitude, while medium AHP (mAHP) remained unchanged.

Together, these findings reveal a cell-type-specific remodeling of intrinsic excitability in CA1, selectively affecting superficial but not deep pyramidal neurons following CRs ablation.

### Alteration in gamma events

To investigate if the changes in synaptic related genes and intrinsic properties would affect the circuit integration and in vivo function of CA1 pyramidal cells, we recorded extracellular electrophysiological activity in the dorsal CA1 region of freely moving mice using 6-shank, 64-channel high-density silicon probes. After the implant (Supp.Fig.3 A-D), mice were introduced to a dimly lit recording room with a square open-field arena (50 cm × 50 cm × 50 cm), where they performed a 20-minute foraging task to collect scattered food pellets while neuronal and positional data were recorded from the CA1 region (Supp.Fig.3 E-H). Locomotion, including running speed, travel distance, and time spent in the center versus peripheral areas, were unaffected by CRs ablation (Supp.Fig.3 I-N).

To assess whether the afferent circuits providing layer-specific inputs to CA1 pyramidal cells were also functionally altered, we next examined local field potentials (LFPs) as a readout of extrinsic connectivity at the circuit level. Prior work shows that gamma events preferentially occur near the peak-to-falling phase of co-existing theta oscillations (theta–gamma coupling) ^18^ ^19^, and that the strength of theta– gamma cross-frequency coupling correlates with memory performance, potentially supporting local information processing under mnemonic demands ^20^ ^21^. Following prior literature, we defined theta as 4–12 Hz, slow gamma as 20–40 Hz, and fast gamma as 40–90 Hz ^22^(Fig. 3 A-D). During theta-associated running, CR;DTA+ mice differed from CR;DTA- in the amplitude of oscillations in both slow gamma (20– 40 Hz) and fast gamma (40–90 Hz): the normalized power of slow and fast gamma was significantly increased in CR;DTA+ mice (Fig. 3 C-D). These differences were not explained by running speed (Supp. Fig. 3I,3M and 3N).

**Figure 3.**
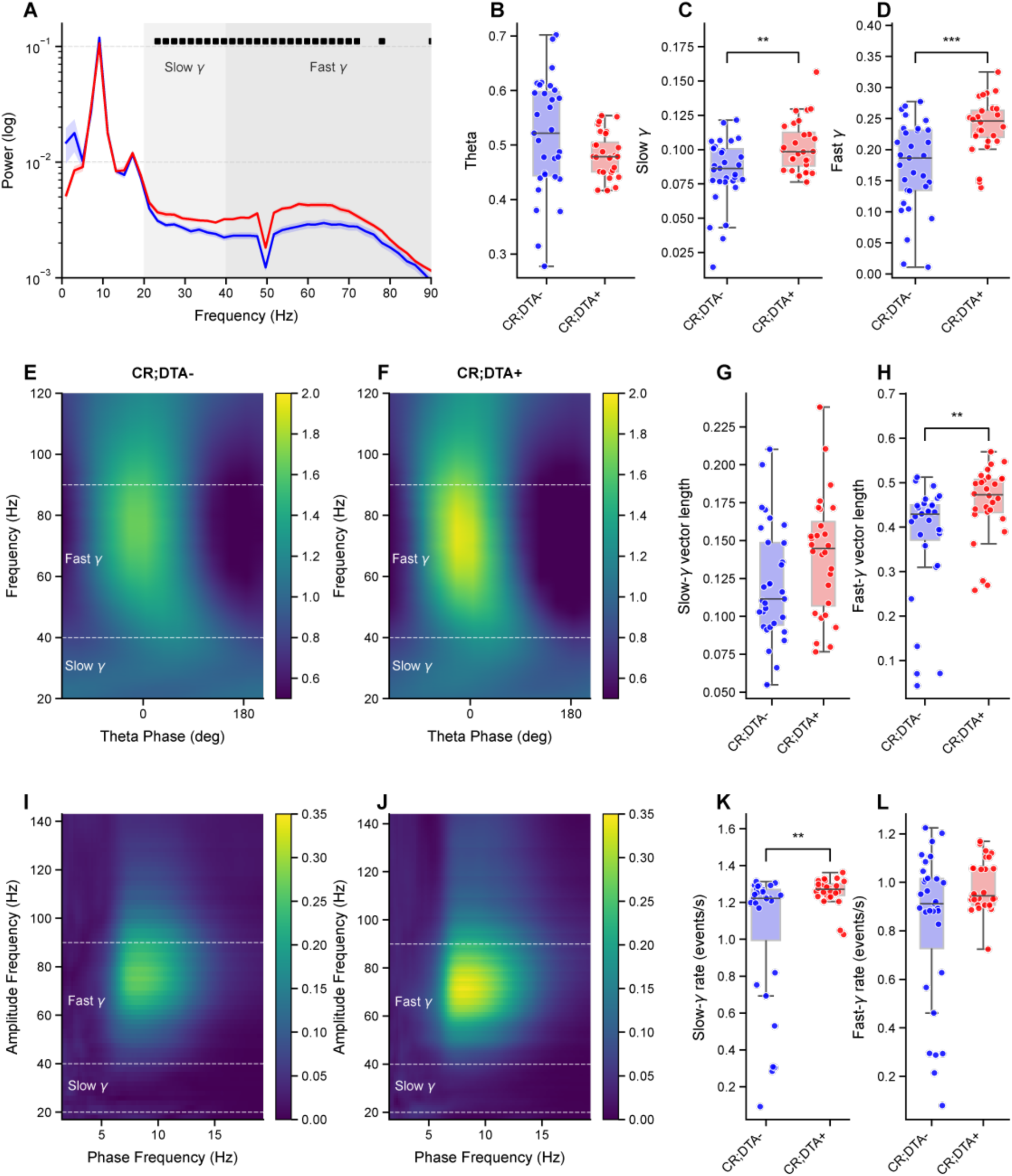
Postnatal ablation of hippocampal CRs increases slow gamma events in CA1. Data are represented as mean ± SEM. Statistical significance was assessed using normal-based or non-parametric tests based on the Shapiro-Wilk test for normality. Normalized power spectra of local field potentials (LFPs) recorded from the pyramidal cell layer during running (>2.5 cm/s), Solid black dots denote significant differences between conditions at specific frequencies (point-wise independent t-tests, Benjamini-Hochberg FDR corrected) Significant differences found between 23.3-151.0 Hz (multiple unpaired t-tests with Benjamini-Hochberg FDR correction, p < 0.05)). (B-D) Power in frequency bands: (B) theta unpaired Student’s t-test, t(57) = 1.678, p = 0.0988 (n_CR;DTA- = 31, n_CR;DTA+ = 28) (C) slow gamma, Mann-Whitney U test, U = 254.0, p = 0.0064 (n_CR;DTA- = 31, n_CR;DTA+ = 28)and (D) fast gamma, unpaired Student’s t-test, t(57) = -4.161, p < 0.001 (n_CR;DTA- = 31, n_CR;DTA+ = 28). (E) Normalized power spectrograms averaged across all theta cycles from CR;DTA- animals. (F) Same as E but from CR;DTA+ animals. (G) vector length of theta–slow gamma coupling. Unpaired Student’s t-test, *t*(57) = -1.910, *p* = 0.0612 (n_CR;DTA- = 31, n_CR;DTA+ = 28). (H) vector length of theta–fast gamma coupling. Mann-Whitney U test, U = 229.0, *p* = 0.0019 (n_CR;DTA- = 31, n_CR;DTA+ = 28). (I) Example cross-frequency coherence plots showing that oscillatory powers at 20–40 Hz and 40–80 Hz frequency (y axis) were modulated by the theta phase (x axis) in CR:DTA- animals. (J) Same as I but from CR;DTA+ animals. (K) The occurrence rate of slow gamma events. Mann-Whitney U test, U = 238.0, *p* = 0.0030 (n_CR;DTA- = 31, n_CR;DTA+ = 28). (L) The occurrence rate of fast gamma events. Mann-Whitney U test, U = 315.0, *p* = 0.0721 (n_CR;DTA- = 31, n_CR;DTA+ = 28). \**p*<0.05*, **p<*0.01.

We quantified the degree of theta–gamma coupling using the phase-locking vector length (Fig. 3 E-H) ^19^ ^22^ and found that theta–slow-gamma coupling was unchanged (Fig. 3 G), but theta–fast-gamma coupling was increased (Fig. 3H). We then quantified the occurrence rates of gamma events during the theta cycles(Fig.3 I-L). The occurrence rate of slow gamma oscillations was increased in CR;DTA+ mice (Fig. 3K), whereas the occurrence rate of fast gamma remained similar between CR;DTA- and CR;DTA+ mice (Fig. 3L). Together, these results indicate overall enhanced gamma oscillations in CA1.

Specifically, the higher number of slow gamma events and stronger theta fast gamma coupling indicate an enhancement of input in the hippocampal CA1 of CR;DTA+ mice. Furthermore, this pattern suggests that CA1 might have a stronger excitatory drive from the CA3→CA1 circuit or indirectly from medial entorhinal cortex layer 2 →CA3/DG as demonstrated through slow gamma enhancement.

### Specific impairments in spatial representation in superficial but not deep pyramidal cells in CA1

Gamma oscillations are driven by coordinated interlaminar dynamics ^19, 23^. Given the widespread alterations in gamma activity alongside the superficial perturbations observed *in vitro* and via snRNAseq, we hypothesized that CR ablation may disrupt specific translaminar circuits. Therefore, we asked how this perturbation differentially impacts deep versus superficial pyramidal cells at the single-cell level. First, we analyzed the intrinsic firing properties of well-isolated single units from both groups. Single unit quality was comparable across groups, with no differences observed in the L-ratio, ISI violation, or signal-to-noise ratio between CR;DTA- and CR;DTA+ mice (Supp.Fig. 4A-D). Based on waveform and firing properties, single units were classified as putative principal neurons or interneurons (Supp.Fig. 4E). No differences were observed in the distribution of these neuronal types between CR;DTA- and CR;DTA+ mice (Supp.Fig. 4F).

Next, we examined the anatomical distribution of cells along the CA1 radial axis and defined the middle of the CA1 pyramidal layer as the recording site where ripples had the largest amplitude (Fig. 4A-B), as previously done in Sharif and colleagues^24^. Using this line as a separator, we divided our recorded CA1 pyramidal cells into deep and superficial (Fig. 4C). No differences were observed in the relative soma position to the CA1 center for deep or superficial cells between CR;DTA- and CR;DTA+ mice (Fig. 4D).

**Figure 4.**
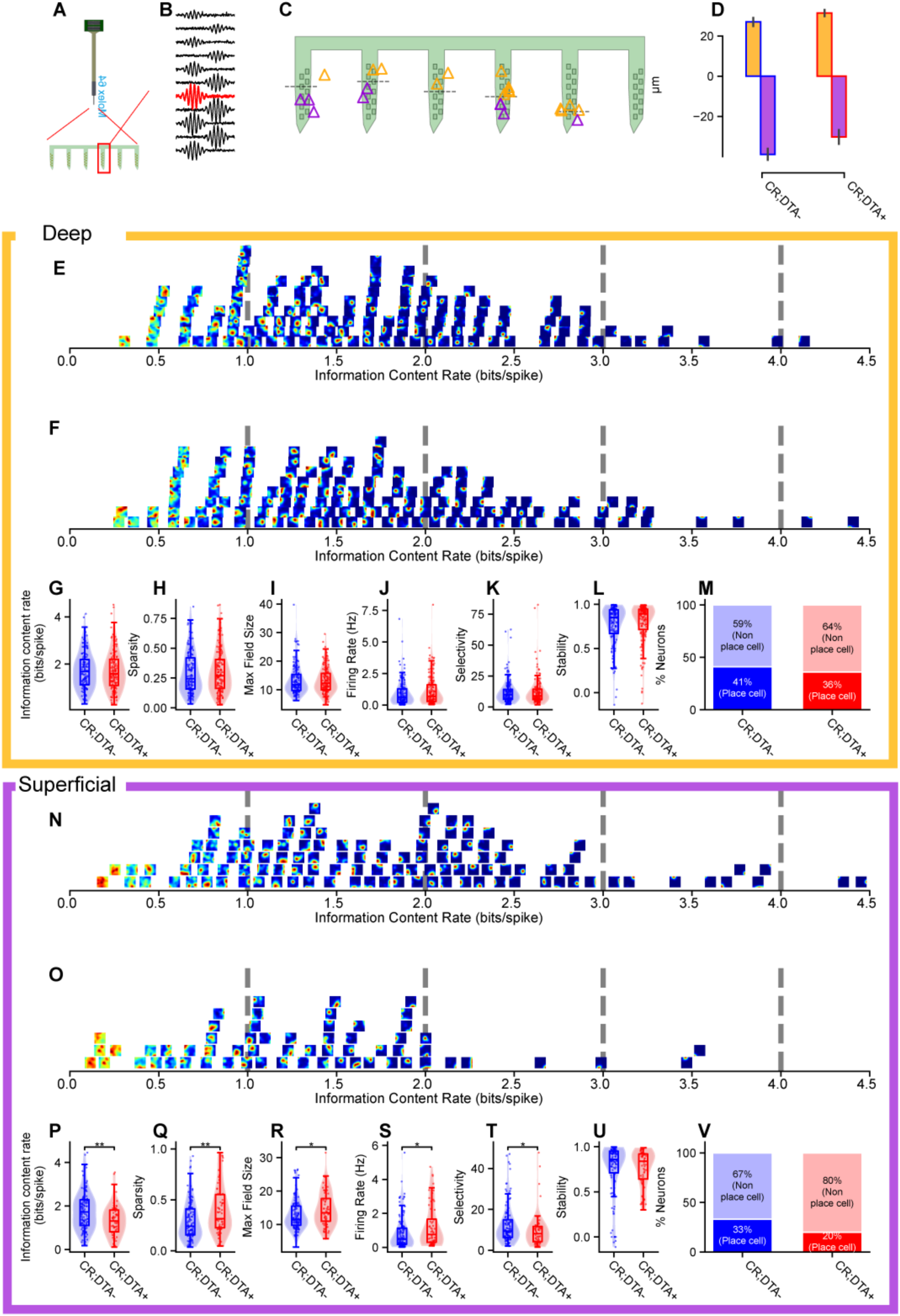
Postnatal ablation of hippocampal CRs more severely impairs spatial representation of CA1 pyramidal cells Data are represented as mean ± SEM. Statistical significance was assessed using normal-based or non-parametric tests based on the Shapiro-Wilk test for normality. (A) Recording schematic: silicon probe targeted to dorsal CA1 with sites spanning the whole CA1 layer. (B) Representative average extracellular local field potential band passed 140 Hz-210Hz for ripple filtering. (C) Example of classified superficial and deep CA1 cell positions relative to the probe layout, with the dashed line representing the center of CA1, identified by the largest amplitude ripple channel. (D) Group summary of the laminar alignment metric (distance to the pyramidal layer, µm) for CR;DTA⁻ (left) and CR;DTA⁺ (right). (E–F) Tile plots of occupancy-normalized spatial rate maps for all putative CA1 pyramidal neurons recorded in CR;DTA⁻ (E, blue) and CR;DTA⁺ (F, red). Tiles are ordered by each unit’s information content rate (spikes/bit; left→right). Vertical dashed guides mark reference values on the information axis. (G–L) Group comparisons (violins with embedded box plots: center line = median; box = 25–75th percentiles; whiskers = 5– 95th; each point = one neuron): (G) Information content rate (spikes/bit). Mann-Whitney U, U = 9474.000, p = 0.808 (CR;DTA- n = 135 cells, CR;DTA+ n = 138 cells). (H) Sparsity (Skaggs sparsity; lower = sparser fields). U = 9059.000, p = 0.6952 (CR;DTA- n = 135 cells, CR;DTA+ n = 138 cells). (I) Maximum field size (percent of spatial bins ≥20% of unit’s peak rate). U = 9164.500, p = 0.818 (CR;DTA- n = 135 cells, CR;DTA+ n = 138 cells). (J) Mean firing rate (Hz). U = 8498.000, p = 0.2106 (CR;DTA- n = 135 cells, CR;DTA+ n = 138 cells). (K) Selectivity (Peak firing rate/ mean firing rate). U = 9637.000, p = 0.6221 (CR;DTA- n = 135 cells, CR;DTA+ n = 138 cells). (L) spatial stability (Pearson correlation between first and second halves of the session). U = 8383.000, p = 0.1532 (CR;DTA- n = 135 cells, CR;DTA+ n = 138 cells). (M) Proportion of place cells per group (stacked bars; place cells vs non–place cells). Place cells are defined by exceeding the shuffle-derived 95th-percentile information threshold. Pearson’s Chi-square test, \\chi^2^ = 0.586, p = 0.4439 (CR;DTA- n = 135 cells, CR;DTA+ n = 138 cells). (N–O) Tile plots as in (E–F) for CR;DTA⁻ (N) and CR;DTA⁺ (O). (P–T, Y) Same metrics as (G–L) for Superficial cells (P) information content rate. U = 4564.000, p = 0.001028 (CR;DTA- n = 115 cells, CR;DTA+ n = 61 cells). (Q) Sparsity. U = 2490.000, p = 0.001569 (CR;DTA- n = 115 cells, CR;DTA+ n = 61 cells). (R) Maximum field size. U = 2817.500, p = 0.03201 (CR;DTA- n = 115 cells, CR;DTA+ n = 61 cells). (S) Mean firing rate. U = 2786.000, p = 0.025 (CR;DTA- n = 115 cells, CR;DTA+ n = 61 cells). (T) Selectivity. U= 4267.000, p = 0.0183 (CR;DTA- n = 115 cells, CR;DTA+ n = 61 cells). (U) Spatial stability. U = 3958.000, p = 0.1618 (CR;DTA- n = 115 cells, CR;DTA+ n = 61 cells). (V) Proportion of place cells per group in Superficial cells. Pearson’s Chi-square test, \\chi^2^ = 2.877, p = 0.08984 (CR;DTA- n = 115 cells, CR;DTA+ n = 61 cells).

When we analyzed spatial representation, we observed that in deep CA1 cells there were no differences in spatial information content, sparsity, max field size, or within-session stability between CR;DTA- and CR;DTA+ animals (Fig. 4G-M), and a similar number of deep cells were classified as place cells. In contrast, among superficial cells, spatial information content (Fig. 4P) and selectivity decreased (Fig. 4T), while sparsity (Fig. 4Q), max field size (Fig. 4R) and firing rate increased (Fig. 4S) compared from control to experimental group. In addition, we observed a trend toward having fewer superficial cells that could be classified as place cells in CR;DTA+ animals (Fig. 4V).

These results demonstrate that, upon early postnatal ablation of hippocampal CRs, the spatial coding remains intact (Fig. 4 L, U), yet spatial representation is more severely impaired in superficial cells. This confirms that the stronger perturbation in superficial pyramidal cell gene expression and intrinsic physiological properties, compared to the deep subpopulation, lead to functional changes in the superficial, and not in the deep, subpopulation of CA1 pyramidal cells.

## Discussion

Our results revealed how deep and superficial pyramidal cells of the CA1 hippocampal subregion are differentially affected by the postnatal ablation of hippocampal CRs. The deep and superficial pyramidal cells in the hippocampal region have historically been treated as homogeneous functional groups in traditional consolidation theory ^25^ ^26^ and in two-stage theory ^27^. However, recent studies report clear differences in their circuit involvement, input-output patterns, and functions ^28–31^ ^32^ ^33^. In particular, spatial representations in deep cells are more strongly anchored to local cues as external reference, whereas superficial cells exhibit coding that is less cue-bound and more internally referenced ^29, 33^. Deep cells are preferentially driven by input from layer 3 of the entorhinal cortex, whereas superficial cells are preferentially driven by internal input from CA3, via the trisynaptic pathway ^28, 34^. In our data, the significant increase in power of gamma oscillations suggests that upstream inputs from entorhinal cortex to CA1 are strengthened following postnatal CR ablation. Interestingly, this enhanced upstream drive differentially impacted CA1 subpopulations. In fact, while the firing activity and spatial representation were never significantly changed among deep cells, in superficial cells the firing rate increased, spikes were more sparsely distributed across space, place-field size was larger, and spatial information content decreased.

At the level of intrinsic excitability, our in-vitro recordings revealed a differential sublayer-specific impact of postnatal CR-cell ablation. Features of deep CA1 pyramidal neurons, were intact between CR;DTA- and CR;DTA+ animals. In contrast, superficial CA1 pyramidal neurons from CR-ablated mice showed clear alterations in intrinsic properties, pointing to a cell-intrinsic re-tuning of spike-generating and integrative properties. Because superficial CA1 neurons are preferentially driven by CA3 input and project back to entorhinal cortex ^28, 34, 35^, increased input resistance and altered fAHPs are expected to boost responsiveness to CA3-driven slow-gamma input while degrading the sparsity and precision of place-related firing. Consequently, even in the presence of enhanced fast-gamma power and prolonged theta-gamma coupling at the network level, the relatively intact intrinsic firing properties of deep cells render them resilient to these circuit-level alterations. Thus, the *in vitro* data provides a mechanistic substrate for the in-vivo phenotype: superficial cells become more excitable but less selective, helping to explain the larger, less informative place fields, reduced selectivity, and impaired spatial representations that we observed explicitly in this sublayer.

Developmentally, the distinction between deep and superficial cells in the hippocampal region reflects both anatomical compartmentalization and birthdate^29, 34–36^. The hippocampal region contains isochronic subpopulations, neurons born together during embryonic neurogenesis that later form distinct functional assemblies with specialized roles in memory consolidation: late-born neurons (superficial pyramidal cells) predominate in short-term recall, while early-born neurons (deep pyramidal cells) are preferentially recruited for long-term memory ^32^. These subpopulations also have a critical postnatal development period where they acquire the differential functionality supporting spatial representation^36^. Our data show that superficial cells are more impaired in gene expression, firing properties and spatial representations. We could therefore speculate about a differential role for postnatal CRs on the maturation of subpopulations of cells born at different time points and following differential postnatal developmental trajectories, with late-born neurons being more dependent on the presence of CRs. Future studies should focus on investigating if memory processes predominantly involving superficial pyramidal cells, such as replaying of awaking activity and memory consolidation, might be specifically impaired by postnatal CR ablation ^34, 35, 37, 38^.

Our single-cell transcriptomics data further highlights the critical role of postnatal CRs in maintaining the transcriptional stability and functional integrity of superficial CA1 pyramidal neurons. The observed reorganization of gene networks likely contributes to the decreased stability and functionality of these neurons, impairing spatial coding and excitability. These observations align with the idea that postnatal CRs are crucial for maintaining the molecular integrity of superficial neurons during development and that their loss may not only lead to immediate functional deficits, as seen in our electrophysiological data, but also prime them for long-term vulnerability in neurodegenerative diseases. This view is consistent with reports that superficial CA1 neurons display greater susceptibility to pathological changes in diseases such as epilepsy and Alzheimer’s disease (AD), where they exhibit earlier neurodegenerative signatures and more severe amyloid and tau pathology ^39, 40^. Moreover, we observed a significant downregulation in the Itm2b network in superficial cells from the CR:DTA+ group. BRI2, encoded by Itm2b, which in certain mutated forms are associated with familial British^41^ and Danish dementias^42^, regulate amyloid-beta (Aβ) protein precursor processing, inhibits Aβ deposition and aggregation, and reduces AD pathology and development^43, 44^. Thus, CRs are essential for the postnatal refinement and stabilization of superficial CA1 neurons, likely providing an important signaling hub to promote connectivity and guide the maturation of CA1 pyramidal cell function. Speculatively, their loss could significantly contribute to the selective vulnerability of this neuronal subpopulation in diseases like epilepsy and AD.

Taken together, our results demonstrate that postnatal CRs are critical for the maturation of hippocampal circuits and function and are preferentially involved in the postnatal development of late-born neurons and the refining of their local network connectivity.

## MATERIALS AND METHODS

### Animals

All experiments were conducted in compliance with protocols approved by the Norwegian Food Safety Authorities and European Directive 2010/63/EU (FOTS ID 24847 and 28607) and performed in accordance with the Norwegian Animal Welfare Act and the European Convention for the Protection of Vertebrate Animals used for Experimental and Other Scientific Purposes. Pups were separated from the mother on postnatal day 21. Adult animals were housed with up to five animals per cage and kept in an inverted 12-hour light/dark cycle, with enriched cages, and food and water ad libitum.

Pde1c-Cre transgenic mice (B6.FVB(Cg)-Tg(Pde1c-Cre) IT146Gsat/Mmucd, MMRRC 030708)^45^ were bred with NDNF-flox-FlpO transgenic mice^46^, kindly donated by Rob Machold, NYU, to create compound heterozygous Pde1c-Cre+/-; NDNF-flox-FlpO+/- mice^5^. Animals of both sexes were used for experiments.

### Viral Injection Procedure

All pups were subjected to viral injections at P0. These were bilateral injections with a recombinant adeno-associated virus (pAAV-mCherry-flp-dtA, modified by the Viral Vector Core facility of the Kavli Institute for Systems Neuroscience, NTNU, from Addgene plasmid #58536; http://n2t.net/addgene:58536; RRID: Addgene_58536) ^47^ expressing Flp-dependent diphtheria toxin A fragment (AAV-DTA). Animals were anesthetized with isoflurane (3%) and head fixed in a stereotaxic frame (Koppf) with a custom-made adaptor. The skin was stretched using standard lab tape with a diamond cutout. Injection coordinates were calculated from *lambda* (AP: +0.8 mm; ML: ±1.2 mm; Z: - 1.22 mm) in each mouse. A virus-filled glass pipette (Drummond Scientific Company) was attached to an injector, Nanoject III (Drummond Scientific Company), used for the injections (48.4 nl, rate 4.8 nl/sec, with 20s postinjection).

Injected Cre^+^ and FlpO^+^ animals (Pde1c-Cre^+/-^; NDNF-flox-FlpO^+/-^; DTA) were considered as the experimental group (CR;DTA+) and their Cre^-^ and FlpO^-^ littermates (Pde1c-Cre^-/-^; NDNF-flox-FlpO^-/-^ ;DTA) were considered as control group (CR;DTA-). Only animals which had cells expressing mCherry in the inner blade of the Dentate Gyrus and dorsal CA1 were used for experiments as this indicated viral spread to most of the hippocampus.

### In-Vitro electrophysiological recordings

CR;DTA+ and CR;DTA- mice of either sex (10-13 weeks old) were anesthetized with isoflurane and euthanized with an overdose pentobarbital (i.p. 100 mg/kg, Apotekerforeninger), followed by intracardial perfusion with cutting solution (4 °C) with (in mM) : 93 choline chloride, 3 KCl, 1.25 NaH2PO4, 30 NaHCO3, 20 HEPES, 10 glucose, 5 MgCl2, .5 CaCl2, 5 N-acetylcysteine, saturated with 95% O2 and 5% CO2. Brains were removed and glued to a vibratome-stage, submerged in cutting solution (4 °C). In the vibratome (VT1000, Leica), acute hippocampal sections (300 µm) were prepared coronally. Sections were bisected, separating the hemispheres, before being transferred to heated cutting solution (34 °C), to recover for 15 min. After recovery, sections were transferred to holding solution (RT), with (in mM): 92 NaCl, 3 KCl, 1.25 NaH2PO4, 30 NaHCO3, 20 HEPES, 10 glucose, 5 MgCl2, .5 CaCl2, 5 N-acetylcysteine, saturated with 95% O2 and 5% CO2.

One control animal, yielding one superficial and one deep cell for the analysis, was prepared slightly different. For this animal, the cutting solution (4 °C) contained (in mM) :87 NaCl, 2.5 KCl, 1.25 NaH2PO4, 26 NaHCO3, 74 sucrose, 10 glucose, 7 MgCl2, 0.5 CaCl2, saturated with 95% O2 and 5% CO2. The same solution was used for heated recovery (34 °C, 15 min), before cooled to RT. All the other steps in preparation and recording were identical.

Sections were recorded from in a heated (∼35°C) recording chamber of an upright microscope (Zeiss Axio Examiner.D1, Carl Zeiss, Jena, Germany) filled with recording solution, with (in mM): 124 NaCl, 3 KCl, 1.25 NaH2PO4, 26 NaHCO3, 10 glucose, 1 MgCl2, 1.6 CaCl2, saturated with 95% O2 and 5% CO2. Sections were held in place using a platinum ring with nylon strings, and hippocampal CA1 was first identified with DIC optics using a 10× (Carl Zeiss, EC Epiplan 10×/0.25 M27) objective. Pyramidal cells with a healthy appearance were chosen based on the position of their soma, close to either the deep or superficial part of the pyramidal layer. All cell selection and recording were performed with a 40× (Carl Zeiss, W N-Achroplan 40×/0.75 M27) objective. Whole-cell patch clamp recordings were done with borosilicate pipettes (Harvard Apparatus, Massachusetts, USA) with a resistance of 3-8 MΩ, filled with an internal solution, with (in mM): 30 K-gluconate, 10 KCl, 10 HEPES, .2 EGTA, 4 ATP-Mg, .3 GTP- Na, 5 phosphocreatine-Na2, pH 7.3. 3% Biocytin was added to the internal solution on the same day as recording. All recordings were done with a Multiclamp 700B amplifier (Molecular Devices), 1550B Digidata (Molecular devices), controlled with Multiclamp commander 700B and Clampex 11 (Molecular Devices). Data were sampled at 20 kHz. All cells were held at -65mV when recording. Protocols and recordings of electrophysiological properties in current clamp mode (as described in Courcelles et al., 2024):

Input resistance (IR, MΩ): measured from Ohm’s law from the peak of voltage responses to -50pA hyperpolarizing current step.

Rheobase (Rheo, pA): First current step to evoke 1-3 action potentials (AP) during a 1-s long depolarizing ramp (500 pA/s). First current step of recording was set below firing threshold and increased with 5 pA per sweep.

Following properties were extracted from a frequency-current (F-I) protocol, with a 1-s long depolarizing step, starting from 0 pA, increasing with 25pA per subsequent step.

Fmin (AP/s): First current step during the F-I recording that evoked at least one AP

Fmax (AP/s): First current step where a cell reached its highest firing frequency.

For following properties all APs were extracted from the sweep which corresponded to first sweep with evoked action potential (AP) + 100 pA:

Fast afterhyperpolarization (fAHP, ΔmV): Measured as difference in mV between AP threshold and the minima within 5 ms after AP. Ap threshold was defined as point where the rise of the AP was 20 mV/ms.

Medium AHP (mAHP, ΔmV): Same as fAHP, but minima located between 5-20 ms after AP.

Depolarization slope (mV/ms): Defined as the maximum derivative of the voltage, after AP threshold, during AP upstroke.

AP amplitude (mV) Defined as peak of AP, measured from AP threshold.

All cells were visualized to confidently locate them in their respective part of the pyramidal layer and to confirm their identity. 9 out of 48 cells were excluded because they were either not in CA1, wrong cell type, unstable recordings, all dendrites cut proximally to soma, or that they could not be recovered with biocytin.

All analysis were performed with python (v3.11.13; Python Software Foundation, 2023), run in an Anaconda environment (v2023.11; Anaconda, Inc., 2023). With scripts ran in Spyder (6.0.7; Spyder Project Contributors 2023), using PyABF (v2.3.8) to read .abf files from patch clamp recordings. Data was statistically tested using Welch’s t-test.

### Histology for in-vitro electrophysiology

After recording, sections were moved to 4% paraformaldehyde, in which they were stored overnight (4 °C). Before immunohistochemical staining, sections were stored in 2% dimethyl sulfoxide (4 °C). For staining, sections were first washed in PBS 3 × 10 min before being incubated in blocking solution containing 10 % normal donkey serum (Jackson ImmunoResearch) and 0.1 % Triton X-100 (Merk) dissolved in PBS, for 1 hour. Next, sections were incubated with primary antibodies in a buffer containing 1 % normal donkey serum and 0.1 % Triton X-100 in PBS, for three overnights (4 °C), while rotating. Primary antibodies: Guinea pig anti-NeuN 1:1000 (Merk, ABN90P), Rat anti-RFP 1:1000 (ChromoTek, AB_2336064). After primary incubation, sections were rinsed 3 × 10 min in PBS, before incubating with secondary antibodies with a similar incubating buffer, overnight at room temperature while rotating. Secondary antibodies: Donkey anti-Guinea pig AF 405 1:500 (Jackson Immuno Research Lab, 706475148, Donkey anti-Rat AF 594 1:500 (Invitrogen, A21209), Streptavidin-conjugated AF633 1:500 (Invitrogen). Before mounting, the sections were washed in PBS 3 × 10 min. They were then mounted on microscope glass slides (Thermo Scientific, Superfrost Plus®, 25×75×1), using electrical tape with a square cut-out to match section thickness. Lastly, glasses were coverslipped (Menzel-Gläser, 15×15 mm, #1) using Fluoromount (Invitrogen, Fluoromount-G) as embedding medium.

### Confocal imaging of thick sections

Single cells were imaged at high-resolution using a Zeiss LSM 990 Airyscan (Carl Zeiss, Jena, Germany). All confocal images were acquired with a Plan-Apochromat 20x/0.8 M27 objective. The pinhole size was set to 1 AU, bit depth: 8 bit, and laser intensity and gain was tuned to be within dynamic range. Additionally, pixel dwell was set to shortest duration and averaging at 1. Files were saved as .czi before being transferred.

### Perfusion and section preparation

Animals were first anesthetized with 0.1 ml isoflurane in a closed chamber and subsequently injected with a lethal dose of pentobarbital (100mg/kg, 0.1ml, IP, SANVIO Pharma AS). Following, the animal was transcardial perfused with ice cold (4 °C) PBS for 5 min (3.5ml / min) followed by freshly made, ice cold, paraformaldehyde (PFA, pH 7.4, Merck Life Sciences AS) for 5 min (3.5ml / min). After recovery of the brain, it was post-fixated in PFA for 16-24h in the fridge. Brains were cryoprotected in increasing concentrations of sucrose (15%, 30%) in preparation for sectioning. brains for cell counting were cut in coronal orientation, on a freezing microtome (Sliding microtome HM 430, Thermo Scientific, Walldorf, Germany) with 50 µm thickness, in three series, before being stored in antifreeze (40% 125 mM phosphate buffer (PBS), 30% glycerol, 30 % ethylene glycol). Brains for silicon probe implanted animals were first embedded in OCT mounting media (VWR international), before being cut coronally, on a cryostat (Cryostar NX70, Thermo Scientific, Walldorf, Germany) with a 30 µm thickness. The sections were immediately mounted on microscope glass slides (Thermo scientific, Superfrost Plus, 25×75×1 mm) and stored at -20 °C.

### Immunohistochemistry

After the recordings were concluded, animals were euthanized and transcardially perfused with phosphate buffered saline (PBS) and subsequently a 4% paraformaldehyde (PFA) solution in PBS (14 ml) at a flow rate of 3.7 ml/min. Brains were dissected, post-fixed in 4% PFA for 3h post-fixation, and transferred to a cryoprotective solution (40% PBS, 30% glycerol and 30% ethylene glycol) for storage at - 20°C. For cryoprotection, brains were briefly rinsed in PBS (10 min) and then transferred sequentially to 15% and 30% sucrose solutions, where they remained for 24 h each at 4 °C. They were sectioned coronally (50 μm) on a freezing microtome (ThermoScientific), collected in three series, and stored in cryoprotective solution at -20°C. For immunohistochemistry, sections were rinsed three times in PBS (10 min each), then incubated for 30 min in a blocking solution containing 10% normal donkey serum (Jackson ImmunoResearch,) and 0.1% Triton X-100 (Merck) in PBS. Primary antibody incubation was performed for three nights at 4 °C with gentle shaking in an incubation solution (1% normal donkey serum and 0.1% Triton X-100 in PBS) containing the following primary antibodies for the CRs counting: rat anti-RFP (1:1000, Chromotek, 5F8), rabbit anti-p73 (1:500, Abcam, ab40658), and goat anti-reelin (1:500, R&D Systems, AF3820), for the silicon probe implanted animal rat anti-RFP (1:1000, Chromotek, 5F8), rabbit anti-p73 (1:500, Abcam, ab40658),. After primary incubation, sections were rinsed three times 1h in PBS and incubated overnight at 4°C with gentle shaking in incubation solution containing the secondary antibodies for the CRs counting: donkey anti-rabbit Alexa 647 dilution: 1:500 (Invitrogen, A31573), donkey anti-rat Alexa 594, dilution: 1:500 (Invitrogen, A21209), and donkey anti-goat Alexa 488, dilution: 1:500 (Invitrogen, A32814). Secondary antibodies for the silicon probe implanted animal: donkey anti-rabbit Alexa 488 dilution: 1:500 (Invitrogen, A-21206), donkey anti-rabbit Alexa 594 dilution: 1:500 (Invitrogen, A-21207). On the following day, sections were washed 3 times in PBS (10 min each), mounted onto SuperFrost glass slides (Thermo Fisher Scientific) in a petri dish containing PBS, rinsed in distilled water for 5 min, and airdried. Slides were coverslipped using Fluormount-G (ThermoFisher), sealed with nail polish, and stored at 4°C.

### Confocal Imaging and Cell Count

The stained sections were imaged using a LSM880 Zeiss Confocal Microscope. Tiled z-stacks were acquired from the dentate gyrus and hippocampal CA regions, including the hippocampal fissure, using a Zeiss Plan-Apochromat 20x/ 0.8 NA objective. Four coronal sections per animal were selected based on strong mCherry/RFP signal, indicating effective viral spread throughout the hippocampus. Sections were spaced 150 μm apart, spanning a dorsoventral distance of at least 600 μm. Image stacks were imported into Neurolucida 360 (Micro Bright Field Bioscience) for analysis. The outer blade of the dentate gyrus and the hippocampal fissure were manually delineated. Cells co-expressing p73 and reelin were marked and counted as CRs. All labeled cells within these regions were counted. Since the sections were non-consecutive, potential overcounting along the z-axis was not considered problematic and no correction was applied. For quantification of the cells, Neurolucida Explorer (Micro Bright Field Bioscience) was used, and the resulting files were exported to Excel. Cell counts were quantified using Neurolucida Explorer and exported to Microsoft Excel. CRs density was calculated as the number of labeled cells divided by the measured area of the delineated region (cells/mm²).

### Scanner imaging

Full series of mounted sections were imaged with a Zeiss AxioscannerZ.1 (Carl Zeiss) controlled with Zen Blue software. All scanner images were acquired with a Plan-Apochromat 20x/0.8 M27 objective, with a resolution of ∼0.33 µm in X and Y, and 4.0 µm in Z. All images were acquired as Z-stacks with four slices across ∼15 µm. Four channels were acquired, AF 405 was excited by led-module 365 nm, with emission filter set at 412-438 nm. AF 488 was excited by led-module 470 nm, with emission filter set at 501-538 nm. AF 594 was excited by led-module white with beam splitter 560, with emission filter set at 570-640. Lastly, AF 647 was excited by led module 625 nm, with emission filter set at 662- 756 nm. All images were saved as .czi format with one image file containing the full microscope glass. The images were later split into single sections.

### Dissection of dorsal hippocampus

Mice were sacrificed at P30 using isoflurane followed by a lethal injection of pentobarbital intraperitoneally administered. Brains were excavated following decapitation into cold, carbogenated ACSF and were promptly processed in cold, carbogenated ACSF on ice under stereotaxic fluorescence guidance. Brains were processed by first dividing hemispheres along the midline. Excess tissue was removed and hippocampi were excavated carefully from ventricles. Fluorescence was used to guide further bisection between dorsal and ventral hippocampus and excess tissue and fibers were cleared away such that only injected tissue (RFP+) was used. Samples were snap frozen in liquid nitrogen and remained in liquid nitrogen in cryosafe vials until processed. Samples were further only included for single nuclei extraction if injection site covered >50% of the dorsal half of hippocampus in both hemispheres.

### Single nuclei RNA extraction

Frozen samples were thawed on ice and processed in an RNAase-free environment using a previously published protocol ^48^. Briefly, tissue was homogenized using cold dounce homogenizers to generate nuclei suspensions, washed 3 times in nuclei-safe buffer, and were then centrifugated at 13,000xg for 45 minutes at 4°C on a sucrose gradient to remove myelin. Isolated nuclei were then adjusted to a concentration of 1000 cells/ul and were sequenced by the NTNU Genomics Core using 10X v3, 3’ barcoding kit and a Nova Seq 6000.

### Single nuclei RNA sequencing analysis

Sequencing data was aligned and demultiplexed using CellRanger followed by cellBender to denoise incorrect barcoding. Count matrices were then analyzed using RStudio and Seurat v5 ^49^. Seurat objects were generated for each sample for a total of 4 samples: 2 CRs; DTA-, 2 CRs; DTA+, each group sex matched. Each sample was annotated for experimental group and sex before performing PCA, clustering, and kNN. Cell clusters were annotated using known markers for subtypes as well as the Allen Brain Atlas. Samples were then integrated using CCA and downstream analyses between CA1 cells for genetic expression changes between CRs; DTA+ and CRs; DTA- were performed at the cell-type and subtype level. Differential Gene Expression (DGE) was performed using MAST, gene expression was plotted using scCustomize RStudio package, and further correlational gene network inferences were made using hdWGCNA^14^.

### Probes and surgery

Anaesthesia was induced by placing the mice in a closed glass induction chamber filled with 3 % isoflurane (air flow: 0.6 l/min; oxygen flow: 0.6 l/min). The mice were given subcutaneous injections of buprenorphine (Temgesic, 0.03 mg/kg), meloxicam (Metacam, 1 mg/kg). The local anaesthetic bupivacaine (Marcain, 1 mg/kg) was injected subcutaneously before the incision was made. After injections the mice were moved to a raised platform with a mask and a stereotactic frame, and the head was secured in place with a set of ear bars. The mice’s body was resting on a heating pad (37°C) to ensure that the body temperature was maintained throughout the surgery. Isoflurane levels were gradually reduced to 0.5-1.5 % (air flow: 0.3 l/min; oxygen flow: 0.3 l/min). Depending on the physiological condition of the mice, which was evaluated by reflex responses and breathing patterns.

Every mouse was implanted with a multi-shank silicon probe attached to a movable nano-drive and equipped with a Cambridge NeuroTech Mini-AMP-64 amplifier (Cambridge NeuroTech). The probes used were ASSY-236 (Cambridge NeuroTech) which coated with Did dye (Thermo Fisher,#D7757). Each ASSY-236 probe consists of six shanks spaced 200 μm apart horizontally, with each shank containing 10–11 recording sites spaced 15 μm vertically and 16.5 μm horizontally. Implant coordinates were determined by two reference points for shank 1 and shank 6 (shank 1: AP 1.8 mm, ML 1.1 mm from bregma; shank 6: AP 2.5 mm, ML 1.8 mm from bregma). A 1.2 mm craniotomy was performed using a Trephines for Micro Drill (FINE SCIENCE TOOLS) to expose both coordinate points, such that shank 1 and shank 6 were aligned with the respective targets and lowered 500–600 μm below the dura. The nano-drive was secured to the skull using an adhesive (OptiBond, Kerr Dental) and dental cement (Meliodent). One screw was placed in the occipital bone over the cerebellum and was connected to the probe grounds/reference. Subsequently, the Mini-AMP-64 was properly positioned next to the nano-drive, the cable and silicon probe were covered with sterilized Vaseline using low temp cautery, and then the whole implant was covered by the dental cement. The total implantation weight was lower than 3 g.

After the surgery, the mice was left in a heated chamber (30 °C) for 30-60 minutes for the immediate recovery phase, after which it was transferred back to the home cage. Additional doses of buprenorphine were administered 8-12 and 24 hours after the first injection. Meloxicam was administered once every 24 hours for as long as was assessed necessary (usually 3-4 days). The mice were left to recover for 3-5 days before the recordings began.

### Behavioral procedures

Recordings typically started 3-5 days after implantation. The nanodrive were turned down in steps of 32-125 μm per day. To determine if the probes were in the CA1 pyramidal cell layer, 1) cells were recorded as the mice explored the empty open field arena and the presence of theta modulation together, 2) ripples during the sleep sessions were used as criteria. When the criteria were met for the first time (at least 3 shanks enter the pyramidal cell layer), the full experimental protocol was started. The silicon probe was subsequently moved down in steps of 32-64 μm per day at the most, and the experiment was repeated several times over the course of 1-2 months (up to 25 recording sessions per animal).

### Behavior and recording sessions

Animal behavior is recorded by a Basler ace2 camera (Basler AG) equipped with far-red filters. The room is illuminated by two Light Bar-45x100-Red (Basler AG) above the arena. Videos are recorded at 50 Hz, and with each frame’s exposure, a TTL signal is sent to the open ephys acquisition board for synchronization of the signal streams.

Neural activity was filtered, amplified, multiplexed, digitized local field acquired through Mini-Amp 64 (Cambridge neuro tech) and recorded through an Open Ephys acquisition board at a 30 kHz sampling rate (0.1-7,500 Hz, x192; 16 bits).

### Behavioral tracking

Videos were analyzed using DeepLabCut^50^ for multi-animal tracking with a custom-trained model. The model was trained on approximately 3,000 frames, incorporating data from all animals involved in this study. However, this model may be overfitted and is likely suitable only for tracking mice in an openfield arena (50 cm × 50 cm). Thirteen keypoints (snout, left ear, right ear, neck, back 1, back 2, back 3, back 4, tail 1, tail 2, tail 3 and tail 4) were labeled for training. The coordinates of the midbrain were used for position analysis.

Position estimates were based on custom trained multi animal DeepLabCut model. Middle point of two is used for the animal’s position. For rate maps in the random foraging task, data were speed-filtered; only epochs with instantaneous running speeds of 2.5 cm/s or more were included.

### Signal preprocessing

The signal processing was completed using *SpikeInterface*. Binary recording files were loaded, and signals were filtered in two ways using the *Spikeinterface.preprocessing* module for spike sorting and local field potential (LFP) analysis.

### EEG Signal Processing and Filtering

The local field potential (LFP) data were extracted from the electroencephalogram (EEG) recordings, restricted to specific time intervals corresponding to the experimental condition (e.g., running periods). The EEG data were sampled at a frequency of 1250 Hz, notch filter at 50Hz was applied to remove the artificial noise. For each recording channel, the raw LFP signal was processed to isolate the frequency components of interest.

Power spectral density (PSD) of the EEG signal was calculated during the wake episode at a sampling frequency of 1250 Hz. The PSD was computed over a single epoch using the periodogram method, which involves applying the discrete Fast Fourier transform (FFT) to the signal, taking the squared absolute value of the resulting complex values, and scaling by the sampling frequency and signal length to yield power per frequency (in Hz). The output is a real-valued representation with frequencies as indices and corresponding PSD values.

### Theta Band Filtering and Phase Computation

To extract theta oscillations, an additional Butterworth bandpass filter was applied to the broadband-filtered LFP signal, targeting the theta frequency range of 6–10 Hz. The filter coefficients were determined using the same sampling frequency of 1250 Hz, and zero-phase filtering was again used to preserve temporal relationships.

The instantaneous phase of the theta rhythm was computed via the Hilbert transform using *scipy.signal* module. The analytic signal was obtained by applying the Hilbert transform to the theta-filtered data, and the phase was extracted as the argument (angle) of this complex-valued signal, yielding phase values in radians ranging from −π to π.

### Detection of oscillation episodes

To extract periods of gamma oscillatory activity in the LFP, we first computed time-varying power within the frequency bands for each recording. Power at each time point was averaged across the frequency range to obtain time-varying estimates of oscillatory power. Time points were collected when the power exceeded 2 SD of the time-averaged power. Time windows, 160 ms in length, were cut around the identified time points. In each 160 ms segment, the maxima of gamma oscillatory amplitude were determined from the gamma bandpass filtered versions of the recordings. Duplicated gamma oscillatory periods, a common consequence of extracting overlapping time windows, were avoided by discarding identical maxima values within a given gamma oscillatory subtype and further requiring that maxima of a given subtype be separated by at least 100 ms. Individual gamma oscillatory windows were finally constructed from the original, non-bandpass filtered recordings as 400 ms long windows centered around the gamma oscillatory amplitude maxima.

### Relationship of gamma to theta phase

Theta phases at the time points associated with gamma maxima (determined as described above) were collected. Theta phases for each gamma oscillatory event were sorted into 30° bins, allowing the phase distribution of each event to be determined. For a given recording, the distributions of gamma oscillations were normalized by dividing the bins by the total number of gamma oscillatory episodes within a given recording. In this analysis, and in all analyses involving oscillation phase, the oscillation peak was defined as 0°.

### Time–frequency representation of power across individual theta cycles

Time-varying power in 2-Hz-wide frequency bands, from 2 Hz to 140 Hz, was obtained for individual theta cycles using the wavelet transform method described above. Time frequency representations for multiple theta cycles recorded from the same site and session within the same animal were then averaged using the theta phase and then normalized using the total average power in the individual 2-Hz bin across the phase (Schomburg et al., 2014).

### Strength of theta-gamma coupling

The theta phase at the time of gamma oscillation maxima occurrence was calculated by bandpass filtering the LFP in the theta range, performing a Hilbert transform on the filtered signal, and then locating theta phase of individual gamma oscillation maxima. Resultant vectors were calculated from the phase distributions of gamma maxima. Lengths of resultant vectors were used as an index for the strength of theta-gamma coupling.

### EEG Signal Processing and Filtering for Spike sorting

For spike sorting, raw signals were band-pass filtered between 300 Hz and 6000 Hz. Subsequently, the *spikeinterface.sorters* module was used to perform automated spike sorting with the *IronClust* algorithm on shanks with up to 11 sites per shank. Automated spike sorting was followed by manual adjustment of clusters using Phy2.

### Spike sorting and well isolated single unit

Using the manually curated data, waveform and quality metrics of the spike data were generated using the *Spikeinterface.SortingAnalyzer* module. Only well-isolated neurons were retained for further analysis. Well-isolated units were defined as those having a median waveform amplitude > 40 µV; L-ratio < 0.05; ISI violation ratios < 0.2; ^51^). Units were further classified into Narrow spike Interneurons, Wide spike Interneurons, and Pyramidal Cells based on their trough-to-peak delay and bursting index ^52, 53^. Briefly, Interneurons are selected by 2 separate criteria: Narrow interneuron assigned if troughToPeak <= 0.425 ms, Wide interneuron assigned if troughToPeak > 0.425 ms and acg_tau_rise > 6 ms. The remaining cells are assigned as Pyramidal cells ^53^. Due to the limited number of cells that were classified in the Wide spike interneurons, they were excluded from further analysis.

### Ripples detection

To detect Ripples, the wide-band signal from CA1 pyramidal cell layers was band-pass filtered (difference-of-Gaussians; zero-lag, linear phase FIR), and instantaneous power was calculated by clipping at 3 SD, rectified, and low-pass filtered. The low-pass filter was at a frequency corresponding to the top cycles of the mean bandpass (for 80-250 Hz band-pass, the low-pass was 55 Hz). Subsequently, the power of the non-clipped signal was computed, and all events exceeding 3 SD from the mean were detected. The events were then expanded until the (non-clipped) power fell below 1 SD; short events (<15 ms) were discarded.

### Classification of deep and superficial CA1 pyramidal cells

To determine the cells’ position relative to deep and superficial CA1, we aligned and averaged all the ripples detected by a shank during the whole recording session and used the channel with the largest signal as a reference for the position of each cell^35^.

### Firing rate maps

Position estimates were convolved with a 20-point Gaussian window and *x*, *y*-coordinates were sorted into 2.5 cm × 2.5-cm bins. Spike timestamps were matched with position timestamps. Only spikes collected at instantaneous running speeds above 2.5 cm/s were included. Firing rate distributions were determined by counting the number of spikes and assessing time spent in 2.5 cm × 2.5-cm bins of the firing rate maps. The distributions were subsequently smoothed with a 2D Gaussian kernel with s.d. of 2 bins.

### Place field area

A firing field in the open field was estimated as a contiguous region of at least 7.5 cm x 7.5 cm where the firing rate was above 20% of the peak rate. Low threshold for peak rate was 1 Hz. Additional fields were identified by deleting the detected field from the rate map and iterating the search for contiguous firing regions in the remaining part of the rate map until no additional fields were found.

### Spatial information score

For each cell, the spatial information score in bits per spike was calculated from the recordings in the open field task as

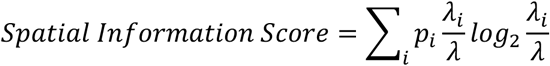

where *λ_i_* is the mean firing rate of a unit in the *i*th bin, λ is the overall mean firing rate, and *p_i_* is the probability of the animal being in the *i*th bin (occupancy in the *i*th bin / total recording time^54^. An adaptive smoothing method, introduced by^55^, was used before the calculation of information scores^56^.

### Sparsity

Spatial sparsity was used to measure how compact and selective the place field of each place cell is relative to the recording enclosure. The spatial sparsity was calculated using the formula as follows^55^:

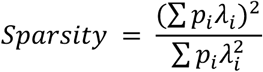

Where pi is the occupancy probability for the animal being at the location of the *i*-th bin in the map and *λ_i_* is the mean firing rate of the cell in bin *i*.

### Stability

The spatial stability within trials was estimated by calculating spatial (2D) Pearson correlation coefficient between firing rate maps generated from the first and second halves of the same trial.

### Defining Place Cells

Place cells were defined as cells with spatial information score above the chance level^57^ among the pyramidal cells. The chance level was determined by a random permutation procedure using all pyramidal cells recorded at that time-point in that region from WT mice. One hundred permutations were performed for each cell in the sample. For each permutation trial, the entire sequence of spikes fired by the cell was time-shifted along the animal’s path by a random interval between 20 s and 20 s less than the total length of the trial (usually 1200 −20 = 1180 s), with the end of the trial wrapped to the beginning to allow for circular displacements. This procedure allowed the temporal firing structure to be retained in the shuffled data at the same time as the spatial structure was lost. Spatial information was then calculated for each shuffled map. The distribution of spatial information values across all 100 permutations of all cells in the sample was computed and finally the 95th percentile was determined. Place cells were defined as cells with spatial information scores above the 95th percentile of the distribution from shuffled data for the relevant group, valid place field and spatial stability > 0.5.

### Euthanasia

For euthanasia, all animals were first anesthetized with isoflurane before being euthanized with a lethal intraperitoneal injection of pentobarbital (100 mg/Kg).

### Blinding Procedures

All experimental procedures, from surgical implantation to manual curation of data, were performed under double-blind conditions by the experimenters to minimize bias where experimenter was not informed of the animals genotype.

### Statistical analysis

Data analyses were performed with custom-written scripts in Python 3.10 and 3.11 (https://www.python.org/d). Open-source Python packages used were: NumPy, SciPy, pandas, and sci-kit learn. Statistical analysis was performed in scipy.stats. Power analysis was not used to determine sample sizes. The study involves two experimental subject groups; therefore, the experimenter was blinded with the animal’s genotype till whole experiments and curation of spike sorting is completed.

## Lead contact

Requests for further information, resources, and reagents should be directed to and will be fulfilled by the lead contact, Giulia Quattrocolo.

## Materials availability

All materials generated in this study are available upon request from the lead contact.

## Acknowledgments

We would like to thank B. A. Zaharia for technical assistance, and H.M. Møllergård for genotyping; R. N. Raveendran and the Viral Vector Core Facility of the Kavli Institute for Systems Neuroscience (NTNU) for viral reagents; P. J. B. Girão for assistance with microscopy; E.Moser, J. Whitlock and M.J.Nigro, for feedback on the manuscript and all the members of the Quattrocolo lab for their insightful comments throughout this project. We further thank the staff in the animal facility at the Kavli Institute for Systems Neuroscience. Special thanks for Dr. H. Nishijo’s valuable comments on data analysis. Single-nuclei RNA sequencing, adaptor trimming and quality control were provided by the Genomics Core Facility (GCF), Norwegian University of Science and Technology (NTNU). GCF is funded by the Faculty of Medicine and Health Sciences at NTNU and Central Norway Regional Health Authority.

The work was supported by a Research Council of Norway (RCN) FRIPRO grants to G.Q. (grant number 324305), RCN Centre of Excellence grants (Centre for Algorithms in the Cortex, grant number 332640; Centre of Neural Computation, grant number 223262; to G.Q.), the Trond Mohn Foundation (Mohn Research Center of the Brain, grant number 2021TMT04, to G.Q.), a NARSAD Young Investigator Grant from the Brain and Behavior Research Foundation (grant number 31551, to G.Q.) and the Kavli Foundation (to G.Q.). The experiments were performed at the NORBRAIN Facility, Norwegian University of Science and Technology (NTNU) (grant number 295721).

## Author contributions

Conceptualization: G.Q., S.

Methodology: S., K.M., K.D., G.Q., N.S., I.L.G.

Software: S.

Data curation: S., K.M., K.D.

Formal analysis: S., K.M., K.D., N.S.

Investigation: S., K.M., K.D., N.S., G.F., I.L.G.

Writing - original draft: S.

Writing - review & editing: S., G.Q., K.M., K.D., N.S., G.F., I.L.G.,

Visualization: S., N.S., K.M., K.D.

Supervision: G.Q.

Funding acquisition: G.Q.

## Data availability

The datasets, source data, and code for reproducing the analyses in this article will be available at the time of publication.

## Supplementary Figures

**Supplementary Figure 1:**
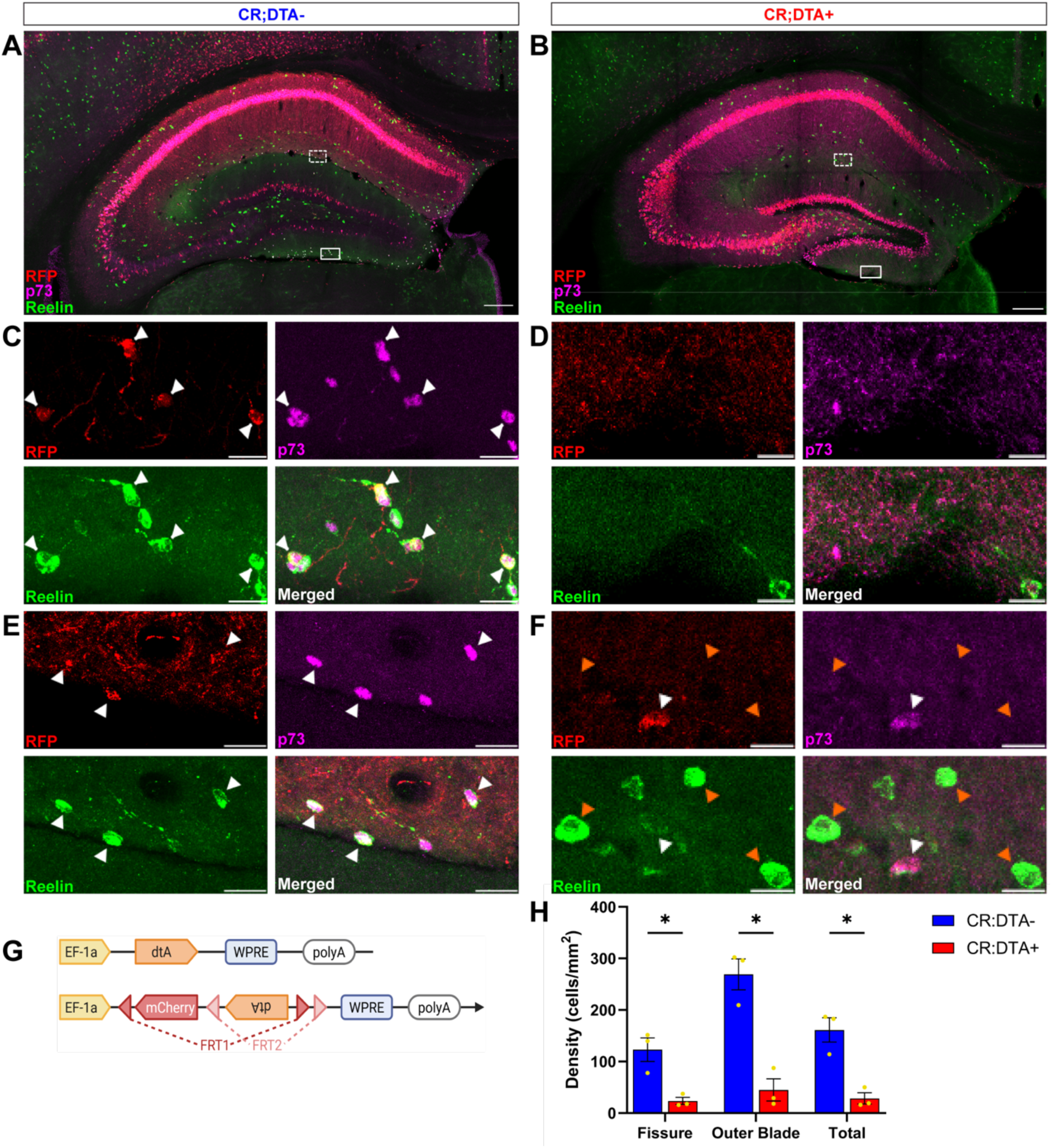
Targeting and ablation of hippocampal Cajal-Retzius cells. Images of coronal hippocampus sections of a (A) CR;DTA- mouse. (B) CR;DTA+ mouse. (A-D) Magnifications of the outer blade of the dentate gyrus outlined (solid line) in A and B, respectively, showing immunolabelled RFP+ (red), p73+ (purple), and reelin+ (green) cells (white arrows); co-localization of p73 and Reelin serves as a marker of Cajal–Retzius (CRs) cells. (E-F) Magnifications of the hippocampal fissure outlined (dashed line) in A and B, respectively, showing immunolabelled RFP+, p73+, and reelin+ cells (white arrows); co-localization of p73 and Reelin serves as a marker of CRs. Cells only positive for reelin (orange arrows; F) are reelin+ interneurons. (G) Schematic of injected viral construct. (H) Comparative analysis of p73+/Reelin+ cell densities between CR;DTA- (n=3) and CR;DTA+ (n=3) animals in the hippocampal fissure, the outer blade of the dentate gyrus and both areas combined (total). Error bars represent standard errors (SE). Statistical significance was determined using Welch’s t-test. The experimental group showed significantly higher values in the fissure (*t*(2.36) = 4.16, *p* = .040), outer blade (*t*(3.63) = 6.08, *p* = .005), and total measurements (*t*(2.82) = 5.12, *p* = .017) compared to the control group. * *p* < .05, ** *p* < .01. Scale bars: 200µm in A and B (overview); 20µm in C-F (representative insets).

**Supplementary Figure 2:**
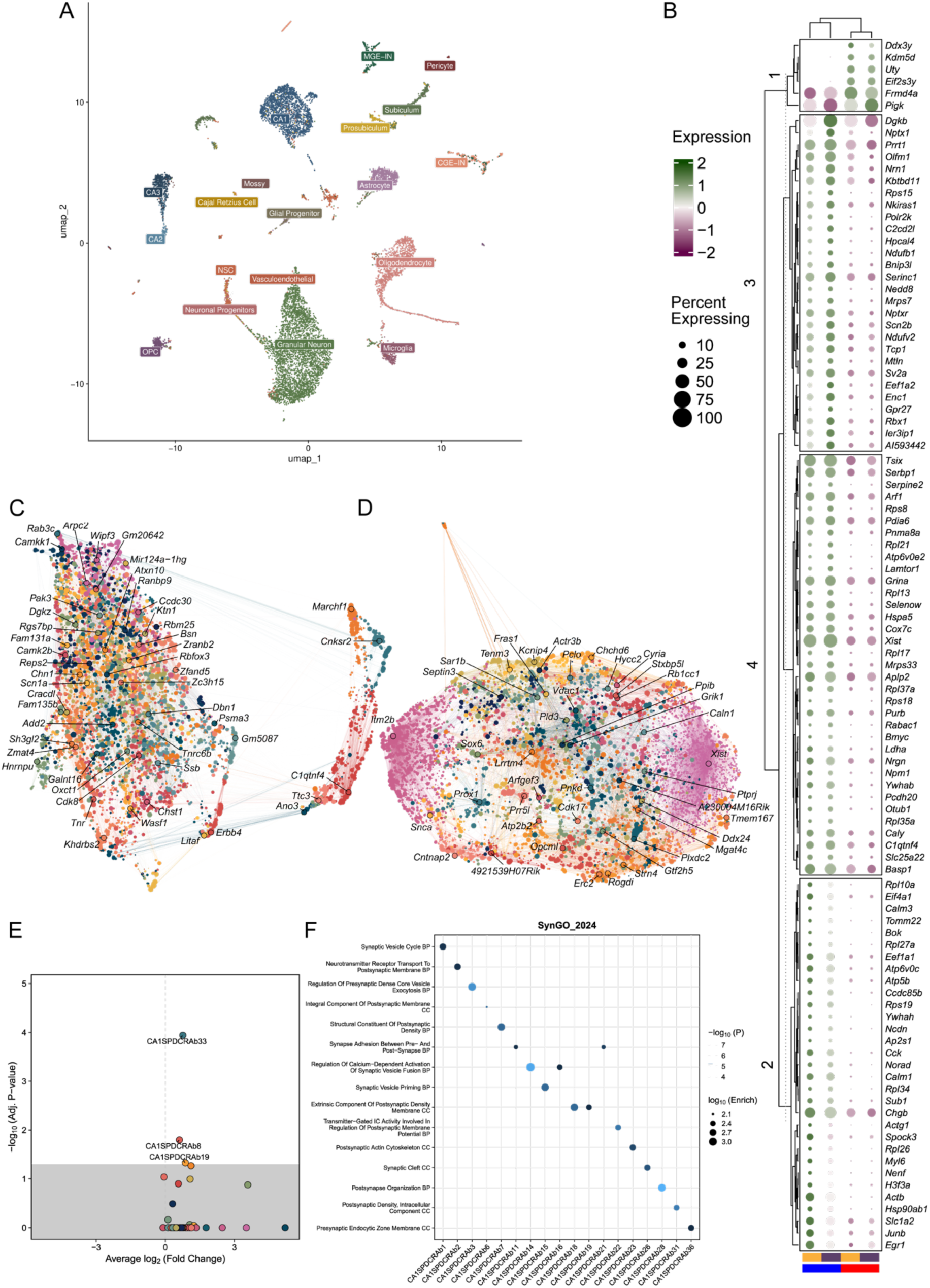
(A) UMAP of snRNAseq data, a total of 11, 619 cells were recovered from dorsal hippocampi of 4 mice. (B) Clustered expression plot of the top 100/637 DEGs between CR;DTA- (blue) and CR;DTA+ (red) CA1 cells. CA1 cells were separated by experimental treatment as well as by superficial or deep subtype identity. K-means clustering was performed on both genes and cell identities (k=4). Gene expression shown is scaled, normalized expression using RNA counts data and constrained to |2|. (C) Gene network UMAP for deep cells in CR;DTA- CA1, demonstrating ∼49 networks of correlated gene expression. (D) Gene network UMAP for deep cells in CR;DTA+ CA1, demonstrating ∼41 networks of correlated gene expression. (E) Volcano plot of differential module eigengene network expression between CR;DTA- and CR;DTA+ CA1 deep layer pyramidal cells. DME was calculated as *Aggregate Network Expression _CRs; DTA+_ – Aggregate Network Expression _CRs; DTA-_*. Cutoff set to adjusted p-val < 0.05 for significantly up or downregulated network. (F) Gene Set Enrichment Analysis of Synaptic Gene Ontology (SynGO) updated 2024 database. Only the top-scoring category is displayed per network. 18/28 networks were found to be significantly enriched, however, none of these networks were significantly up or downregulated in the DME analysis (Supp. Figure E).

**Supplementary Figure 3:**
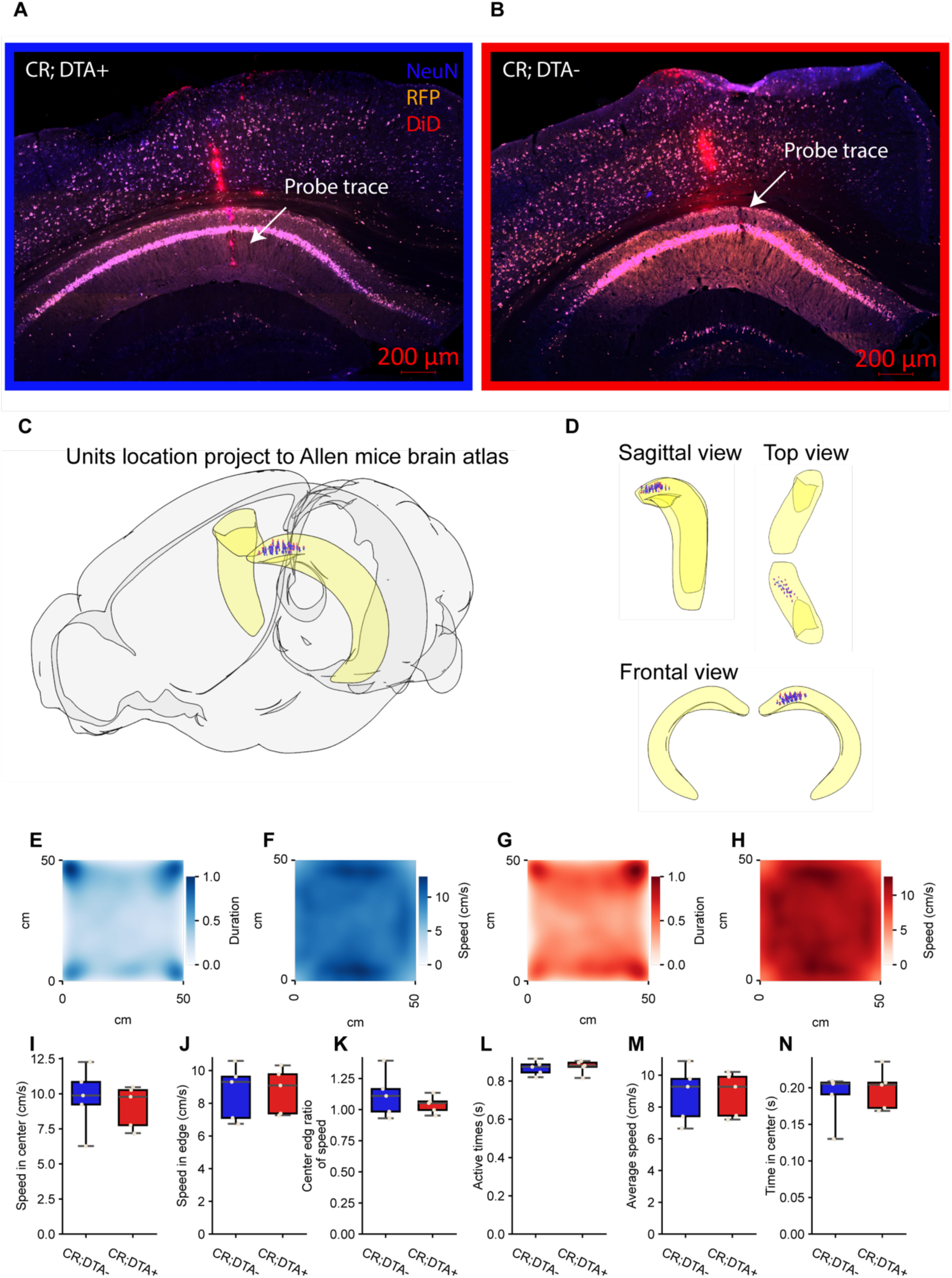
(A) Coronal section of the CR;DTA+ hippocampus stained for RFP (Orange) and NeuN (Blue); DiD (Red) was coated onto the probe before surgery, enabling post hoc reconstruction of the probe track. (B) As in panel A, but from CR;DTA– mice. (C) The reconstructed probe track and unit locations were registered to the Allen Mouse Brain Atlas using QUINT workflow^58^, with the hippocampus shown in light yellow. (D) Alternative view of unit locations within the hippocampus, showing that unit positions are concentrated in the dorsal proximal CA1. (E–H) Heatmaps showing the movement data of CR; DTA− (blue) and CR; DTA+ (red) mice during the exploration task. (A and C) Duration in each location, (B and D) Speed distribution, during the exploration task. (I–N) Behavioral measures of locomotor activity: (I) Speed in the center, (J) Speed in the edge, (K) Center-to-edge speed ratio, (L) Active times, (M) Average speed, and (N) Time spent in the center. No significant differences were observed between CR;DTA− and CR;DTA+ mice in any of the measures (p > 0.05 for all comparisons). Data are represented as median with interquartile range (IQR), and whiskers showing minimum and maximum values. Individual data points represent individual mice (Control n=5, Experimental n=5). Statistical significance was determined using an unpaired two-tailed Student’s t-test. Significance levels: * p < 0.05, ** p < 0.01, *** p < 0.001, **** p < 0.0001, ns = not significant. Exact p-values: (E) p = 0.6299; (F) p = 0.9265; (G) p = 0.4046; (H) p = 0.8097; (I) p = 0.9903; (J) p = 0.6513.

**Supplementary Figure 4:**
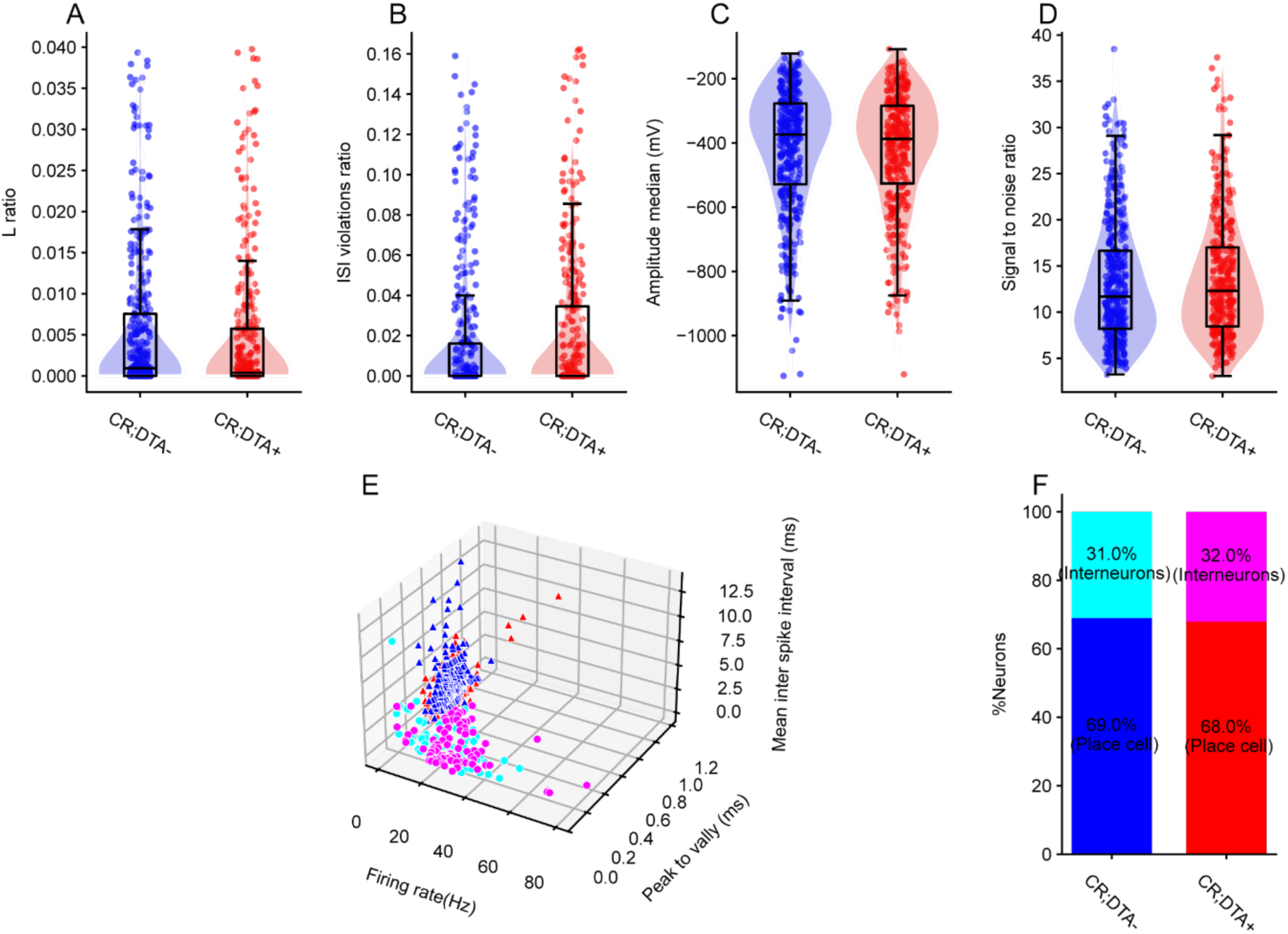
(A) Distribution of cluster isolation quality (L-ratio) for all recorded units in control (CR;DTA−, blue) and CR-ablated (CR;DTA+, red) mice. `Mann-Whitney U test, U = 64708.00, p = 0.0813, n(control) = 378 units, n(experimental) = 318 units.` (B) Inter-spike interval (ISI) violation ratio, reflecting refractory period violations for each unit.`U = 57697.00, p = 0.3157.` (C) Median spike amplitude (µV) across units. `U = 61831.50, p = 0.5129.` (D) Signal-to-noise ratio (SNR) of extracted spikes. `U = 57877.00, p = 0.3999.` (E) Three-dimensional scatter plot of all well-isolated units recorded in CA1 from CR; DTA− and CR; DTA+ mice. Each point represents a single unit positioned by its mean firing rate (x-axis), spike peak-to-valley duration (y-axis), and mean inter-spike interval (z-axis). Units with narrow spike width and high firing rate were classified as putative interneurons (triangles), whereas units with broader spikes and lower firing rates were classified as putative pyramidal cells (circles). Colors indicate experimental group (CR; DTA− vs CR; DTA+). (F) Proportion of putative pyramidal cells and interneurons in CR; DTA− (left bar) and CR; DTA+ (right bar) mice. Stacked bars show that the relative fraction of pyramidal cells (≈69% vs 68%) and interneurons (≈31% vs 32%) is similar between groups, indicating that postnatal CR-cell ablation does not bias the types of neurons recorded. Chi-square test, chi2 = 0.03, p = 0.8599, total n = 656 units.

